# Somatic mutations of activated signaling genes, transcription factors, or tumor suppressors are a precondition for leukemic transformation from myelodysplastic syndromes: a sequencing analysis of 64 paired samples

**DOI:** 10.1101/2021.05.28.446246

**Authors:** Xiao Li, Chun-Kang Chang, Feng Xu, Lin-Yun Wu, Juan Guo, Lu-Xi Song, Yin Tao, Qi He, Zheng Zhang, Dong Wu, Li-Yu Zhou, Chao Xiao, Ji-Ying Su

## Abstract

The transformation biology of secondary AML from MDS is still not fully understood. Here, we performed a large cohort of paired sequences including target, whole-exome and single cell sequencing to search AML transformation-related mutations (TRM). The results showed that fifty-five out of the 64 (85.9%) patients presented presumptive TRM involving activated signaling, transcription factors, or tumor suppressors. Most of TRM (63.6%, 35 cases) emerged at the leukemia transformation point. All five of the remaining nine patients analyzed by paired whole exome sequencing showed TRM which are not included in the reference targets. Single-cell sequencing indicated that the activated cell signaling route was related to TRM which take place prior to phenotypic development. Of note, defined TRM was limited to a small set of genes (less than ten, in the order: NRAS/KRAS, CEBPA, TP53, FLT3, RUNX1, CBL, PTPN11 and WT1, accounted for 91.0% of the mutations). In conclusion, somatic mutations involving in activated signaling, transcription factors, or tumor suppressors appeared to be a precondition for AML transformation from myelodysplastic syndromes. The TRM may be considered as new therapy targets.

---

In most cases, *de novo* acute myeloid leukemia (AML) shows rapid onset without an obvious pre-AML period [1]. Patients have been reported to show genetic abnormalities, such as PML/RARa in the M3 subset [2]; ETO fusion gene in the M2b subset [3]; and CBFβ-MYH11 in M4 EO [4]. In addition, somatic mutations involving CEBPa, FLT3, NPM1, and c-Kit have been used to assess prognosis [5]. Based on these tumor-derived biological characteristics, target-specific and immunological methods have been developed to treat AML [6]. Unlike *de novo* AML, secondary AML (sAML), transformed from myelodysplastic syndromes (MDS), shows unique genetic features. First, there is often an obvious pre-AML stage, where somatic gene mutations result in initial events (clonal hematopoiesis) and/or driver events (development of MDS phenotypes), largely involving epigenetic regulation, RNA splicing, fewer transcription, and others [7,8]. Second, despite preliminary findings on the roles of late-stage gene mutations involving various signaling pathways or transcription during sAML development [9,10], including our primary consideration on sAML-related mutations [11], the transformation biology of sAML is still not fully understood. There are many unanswered questions, as: Could early or late somatic mutations solely involved in epigenetic regulation or RNA splicing also induce sAML? In other word, if it is necessary for sAML transformation by occurrence of some more aggressive gene mutations? Which mutations are the key factors that induce the transformation process? When do the transformation-related mutations emerge? Are they pre-existing at MDS diagnosis, or do they emerge when the transformation starts? Because of the lack of conclusive finding on sAML transformation, we tentatively named those suspected last events as sAML transformation-related mutations (TRM). Hypothesizing that mutations involving signaling pathways, myeloid transcription factors, or tumor suppressors at MDS diagnosis and AML transformation are both a precondition for MDS to progress to sAML, we proposed that analyzing a large cohort of paired samples would help answering to these questions. Examination of over 1,000 MDS patients over 10 years, we identified 64 patients who underwent AML transformation. Paired samples were acquired at MDS diagnosis and immediately after AML transformation. Sequencing for 39 target genes in all samples was followed by exome sequencing in the few patients whose target sequencing did not identify novel mutations. An additional sample pair was subjected to single-cell RNA transcription sequencing. Using these techniques, we obtained data that informs the biology of sAML transformation and suggests novel strategies to prevent leukemic transformation of MDS.

## Patients and Methods

### Sample collection

From January 2004 until October 2019, a total of 1,427 patients were diagnosed with MDS in our department. Over 90% were followed up by calls or with face-to-face consultations, and over 800 patients consented to provide samples for next generation sequencing. DNA was extracted from bone marrow (BM) samples obtained from patients before 2009 and retrospectively used for target sequencing or whole exome sequencing. For patients whose BM samples were acquired in or after 2009, DNA was extracted from the samples and immediately used for targeted sequencing or whole exome sequencing. For single-cell RNA transcription sequencing, fresh BM was used to isolate mononuclear cells within six hours of BM aspiration. Targeted sequencing was performed using paired samples (at the diagnosis and transformation points) to detect transformation-related gene mutations (TRM). Whole exome sequencing was performed when conclusive results for last mutation events were not obtained via targeted sequencing and when adequate residual DNA extracts were available. Single-cell RNA transcription sequencing was used to examine the underlying transformation dynamics. Diagnoses for MDS and sAML were established in strict accordance with the WHO criteria, and the CMML subset was diagnosed according to the FAB classification [12,13].

### Genomic DNA preparation, target enrichment, and sequencing

Genomic DNA (gDNA) was extracted using the DNeasy Blood and Tissue Kit (Qiagen, Germany) according to the manufacturer’s protocol. Genomic DNA was sheared using the Covaris^®^ system (Covaris, USA), and the DNA sample was prepared using the Truseq DNA Sample preparation Kit (Illumina, USA) according to the manufacturer’s protocol.

For probe design, both coding and regulatory regions of target genes were included in the custom panel. The regulatory regions comprised promoter regions (defined as 2 kb upstream of the transcription start site), 5′ un-translated region (5′-UTR), and intron-exon boundaries (50 bp). Custom capture oligos were designed using SureDesign website of Agilent Technologies (Agilent, USA). Hybridization reactions were carried out on ABI 2720 Thermal Cycler (Life Technologies, USA) with the following hybridization conditions. Hybridization mixture was incubated for 16 or 24 h at 65°C with a heated lid at 105°C. After the hybridization reactions, the hybridization mixture was captured and washed with magnetic beads (Invitrogen, USA) and SureSelect target enrichment kit (Agilent, USA). The captured product was enriched with the following cycling conditions, 98°C for 30s, 10 cycles of 98°C for 10 s, 60°C for 30 s, 72°C for 30 s, and 72°C for 5 min. Library quality was assessed using an Agilent 2100 Bioanalyzer (Agilent, USA), and multiplexed sequencing was performed on HiSeq 2500 sequencers with 2×150 paired-end modules (Illumina, USA). Average sequencing depth was 800×. Following 39 sequenced target genes were included: *ANKRD11, ASXL1, BCOR, CALR, CBL, CEBPA, DHX9, DNMT3A, ETV6, EZH2, FLT3, GATA2, IDH1, IDH2, ITIH3, JAK2, KIF20B, KIT, KRAS, MPL, NF1, NPM1, NRAS, PHF6, PTPN11, PTPRD, ROBO1, ROBO2, RUNX1, SETBP1, SF3B1, SRSF2, STAG2, TET2, TP53, U2AF1, UPF3A, WT1*, and *ZRSR2*.

### Whole-exome sequencing

The gDNA library was prepared using a TruSeq DNA Sample Preparation Kit (Illumina, San Diego, CA, US) in accordance with the manufacturer’s protocol. In-solution exome enrichment was performed using a TruSeq Exome Enrichment kit (Illumina) according to the manufacturer’s instructions. The enriched DNA samples were sequenced via 2×100 paired-end sequencing using a Hiseq2000 Sequencing System (Illumina). Illumina Sequencing Control v2.8, Illumina Off-Line Basecaller v1.8, and Illumina Consensus Assessment of Sequence and Variation v1.8 software were used to produce 100-base pair (bp) sequence reads.

### Sequencing data processing, variant calling, and annotation

Before variant calling, the raw sequence reads were mapped to the reference genome (hg19), duplicate reads were marked and removed to mitigate biases introduced by amplification, and base quality scores were recalibrated using the Genome Analysis Toolkit (GATK). The Ensembl VEP and vcf2maf tools were applied to generate a MAF format for somatic mutation annotation, and ANNOVAR tool was used to annotate frequency information of variations in the population database. The variants were identified as low-frequency functional mutation if they had < 0.1 frequency in ExAC03 database, < 0.01 frequency in 1,000 genome database, and < 0.05 frequency in GeneskyDB database. According to the results, the variant was then extracted as one of the following functional annotations: “Frame_Shift_Del/ins”, “In_Frame_ Del/Ins”, “Missense_Mutation”, “Nonsense_Mutation”, “Nonstop_Mutation”, “Splice_Site” or “Translation_Start_Site”.

### Single-cell RNA sequencing and bioinformatic data analysis

Bone marrow mononuclear cells(BMMC) concentrations were measured by hemocytometer and adjusted to 700-2,000 cells/μL. Single-cell RNA-seq libraries were generated using 10x Genomics Single Cell 3’ reagent v2 according to the manufacturer’s instructions. Twelve and 14 cycles were used for complementary DNA and index PCR amplifications, respectively. Amplified cDNA and library quality was assessed using Agilent 2100 Bioanalyzer (Agilent, USA). Libraries were pooled and sequenced on HiSeqX Ten platform (Illumina) with 2×150 paired-end modules generating at least 50K mean reads per cell. Raw sequencing data were processed using Cell Ranger version 3.1.0 (10x Genomics) with default parameters. Quality control metrics were used to remove cells with mitochondrial gene percentage more than 20% or cells with fewer than 200 genes detected. Variably expressed gene selection, dimensionality reduction, and clustering were performed using the Seurat package version 3.1.3. Principal component analysis was performed on significantly variable genes and the first 40 principal components were used for UMAP dimension reduction. Cell type of each cluster was identified using singleR version 1.0.1. For paired samples, differentially expressed genes (fold change > 2 and Wilcox test P value < 0.05) were identified in each of seven cell types independently, of which the mutation genes PTPN11 and NRAS were used for GO and KEGG enrichment analysis.

### Definition of the presumed transformation-related mutation

When we analyzed the paired data, if pre-existing genes at MDS diagnosis or newly-emerging genes at sAML transformation met following conditions, we considered they are sAML transformation related mutations: 1. they must be involved in at least one of the three function pathways namely, active signaling, myeloid transcription or tumor suppression. 2. They emerged after sAML transformation (better weight) or pre-existed at MDS diagnosis (poorer weight). Newly emerged mutations were preferentially considered as the presumed transformation-related mutations. 3. When ≥ 2 suspicious mutations co-existed to be defined as transformation-related mutations, the biologically more aggressive one (active signaling > myeloid transcription > tumor suppressor); or with lower VAF (Various allele frequency) among newly emerged mutations (meaning latest emergence) were defined as transformation-related mutations (TRM).

### Statistics analysis

Statistical analyses were conducted using SPSS software version 18.0. Kaplan-Meier analysis was used to evaluate the time to survival and time to progression. All P-values were based on 2-sided tests and P-values less than 0.05 were considered statistically significant.

## Results

### Target sequencing

Paired samples acquired from 64 MDS patients before and after transformation to sAML were analyzed using target sequencing. Identified somatic mutations are presented in Table 1.

**Table.1.** The results of target sequencing from paired samples of 64 patients.

| No<br>( UPN ) | Diagnosis | Duration<br>( MDS to<br>AML)(m) | Presumed last event | All gene mutations |
| --- | --- | --- | --- | --- |
| ***The patients in whom AML transformation-related mutations could be presumed by paired sequencing analysis***<br>***(listed as activated signaling, transcription factors and tumor suppressor***<br>Activated signaling |  |  |  |  |
| 1.(2998) | RAEB2 |  | RUNX1/9/967+1>A6/886+1>A(31)<br>KRAS/2/G12S(30) | ASXL1;KRAS;RUNX1;U2AF1 |
|  | AML | 2 | RUNX1/9/967+1>A6/886+1>A(28)<br>**KRAS/2/G12S(27)** | ASXL1;KRAS;RUNX1;U2AF1 |
| 2.(3430) | CMML1 |  | GATA2/4/G327E(29)/4/323_323del(13)<br>RUNX1/6/D198Y(41)<br>KRAS/2/G12C(11)<br>NRAS/2/G12D(29) | GATA2*2; KRAS;NRAS;RUNX1;SF3B1;UPF3A |
|  | AML | 7 | GATA2/4/G327E(29)/4/323_323del(38)<br>RUNX1/6/D198Y(43)<br>**KRAS/2/G12C(45)** | GATA2*2; KRAS; RUNX1;SF3B1;UPF3A; EZH2 |
| 3. ( 2356 ) | RAEB1 |  | PHF6/4/C107fs(16)<br>RUNX1/5/A149_A150delinsGX ( 16 )<br>TP53/3/c.74+14T>C(51) | BCOR;ITIH3;PHF6;RUNX1;TET2;TP53 |
|  | AML | 6 | PHF6/4/C107fs(23) | BCOR;ITIH3;PHF6;RUNX1;TET2;TP53 ; EZH2; KRAS |
|  |  |  | RUNX1/5/A149_A150delinsGX ( 21 )<br>TP53/3/c.74+14T>C(55)<br>**KRAS/2/G13D(25)** |  |
| 4. ( 3541 ) | RAEB2 |  | GATA2/5/L345W(43) | DNMT3A;GATA2;TET2 |
|  | AML | 14 | GATA2/5/L345W(30)<br>**KRAS/2/G12R ( 30 )** | DNMT3A;GATA2;TET2;BCOR;KRAS |
| 5. ( 809 ) | RCMD |  | CEBPA/1/K352fs ( 15 )<br>FLT3/9/D358V(48)<br>RUNX1/6/508+1G>C(24)/5/G170R(23) | ASXL1;CEBPA;EZH2;FLT3;RUNX1*2;STAG2;TET2 |
|  | AML | 17 | FLT3/9/D358V(48)<br>RUNX1/6/508+1G>C(45)/5/G170R(42)<br>**NRAS/2G13R(48)** | ASXL1; EZH2;FLT3;RUNX1×2;STAG2;TET2;BCOR; NRAS |
| 6. ( 1057 ) | RAEB1 |  | PTPRD/10/A539V ( 51 ) | DNMT3A; PTPRD;STAG2;U2AF1 |
|  | AML | 48 | PTPRD/10/A539V ( 46 )<br>RUNX1/4/L98fs(28)<br>**NRAS/2/3/G12V/Q61L(15)(15)**<br>**CBL/8/C384R(12)** | DNMT3A; PTPRD;STAG2;U2AF1;CBL;NRAS×2;RUNX1 |
| 7. ( 3435 ) | RCMD |  | RUNX1/1/P86S(46) | ASXL1;EZH2;RUNX1;STAG2;UPF3A |
|  | AML | 33 | RUNX1/1/P86S(43)<br>PTPN11/3/F71L(34) | ASXL1;EZH2;RUNX1;STAG2;;UPF3A;BCOR;NRAS;PTPN11 |
|  |  |  | **NRAS/2/G12D(9.7)** |  |
| 8..(2747) | RAEB2 |  |  | BCOR; ITIH3 |
|  | AML | 10 | **NRAS/2/G12D(57)** | BCOR; ITIH3;KIF20B;NRAS;STAG2; |
| 9 ( 4603 ) | RCMD |  | RUNX1/1/H78Y(53)<br>ETV6/6/R339I(15) | DNMT3A;ETV6;EZH2;KIF20B;RUNX1;SETBP1 |
|  | AML | 5 | RUNX1/1/H78Y(43 )<br>**NRAS/2/G12A(23)**<br>**PTPN11/3/A72V(18)** | DNMT3A; KIF20B;RUNX1;SETBP1;NRAS;PTPN11; |
| 10.( 3626 ) | RCMD |  |  | ASXL1;DHX9;IDH1;RUNX1;TET2;U2AF1 |
|  | AML | 29 | ETV6/6/G381fs(48)<br>**NRAS/2/G12A(44)** | ASXL1;DHX9;IDH1;RUNX1;TET2;U2AF1;ETV6;NRAS |
| 11.( 4687 ) | RCMD |  | GATA2/3/M223I(51)<br>RUNX1/2/G138fs(42) | ASXL1;EZH2;GATA2;RUNX1;STAG2;TET2*2 |
|  | AML | 6 | GATA2/3/M223I(55)<br>RUNX1/2/G138fs(50)<br>**NRAS/2/G12D(52)** | ASXL1;EZH2;GATA2;RUNX1;STAG2;TET2*2;ROBO1;NRAS |
| 12.(1587) | RAEB1 |  | JAK2/14/V212F(46)<br>FLT3/19/c.2208-14A.G(52) | IDH2;FLT3;JAK2;U2AF1;ZRSR2 |
|  | AML | 25 | JAK2/14/V212F(43)<br>**FLT3/19/c.2208-14A.G(49)** | IDH2;FLT3;JAK2;U2AF1;ZRSR2 |
| 13.(2570) | RAEB1 |  | RUNX2/6/D198N(35) | IDH2;RUNX1;SRSF2 |
|  | AML | 5 | **FLT3/11/A445V(49)/11/A445S(49)** | FLT3*2; |
| 14(1792) | RAEB1 |  | RUNX1/9/Q397R ( 47 ) | RUNX1;U2AF1 |
|  | AML | 8 | RUNX1/9/Q397R ( 30 )<br>**FLT3/11//E444V(37)11/E444X)(37)** | RUNX1;U2AF1; FLT3*2;ROBO1; |
| 15.(2117) | RCMD |  | TP53/5/8/Y126C(16)/R273S(15) | SRSF2;TP53*2 |
|  | AML | 8 | TP53/5/8/Y126C(14)/R273S(14)<br>**FLT3/9/D358V(55)** | TP53*2;FLT3 |
| 16.(2577) | CMML1 |  | CEBPA/1/p69-70del(17)<br>NPM1/10/L258fs(33) | CEBPA;DNMT3A;NPM1;ROBO1;TET2 |
|  | AML | 11 | CEBPA/1/p69-70del(21)<br>NPM1/10/L258fs(25)<br>**FLT3/14/K602delinsREYEYDLK (14)** | CEBPA;DNMT3A;NPM1;ROBO1;TET2;FLT3 |
| 17(3183) | RCMD |  |  | BCOR;DHX9;EZH2*2;IDH1;;ROBO1;STAG2;U2AF1 |
|  | AML | 6 | **FLT3/9/D358V(52)**<br>**CBL/11/C1564-13C>T(40)** | BCOR;DHX9;EZH2*2;IDH1;ROBO1;STAG2;;U2AF1;CBL;FLT3;TET2 |
| 18 (2206) | RCMD |  | NF1/17/M645V(49)<br>PTPRD/29A1047S(44)<br>CBL/8/C396R(27 ) | ANKRD11;ASXL1;CBL;NF1;PTPRD;U2AF1 |
|  | AML | 3 | NF1/17/M645V(51)<br>PTPRD/29A1047S(40)<br>**CBL/8/C396R(21)** | ANKRD11;ASXL1;CBL;NF1;PTPRD;U2AF1 |
| 19.(4559) | RAEB2 |  | CBL/15/S791fs(9.6)/15/L790fs(9.7) | ANKRD11;ASXL1;CBL*2; ROBO1 |
|  | AML | 11 | **CBL/15/S791fs(8.7)/15/L790fs(8.8)** | ANKRD11;ASXL1;CBL*2; ROBO1 |
| 20.(692) | CMML2 |  | GATA2/3/S186T(47) | GATA2;ITIH3;PHF6;RUNX1;SF3B1; |
|  |  |  | PHF6/9/:c.968+1G>A ( 25 )<br>RUNX1/9/F416fs(27) |  |
|  | AML | 10 | GATA2/3/S186T(50)<br>PHF6/9/:c.968+1G>A ( 26 )<br>RUNX1/9/F416fs(24)<br>**CBL/9/R420Q(34)** | GATA2;ITIH3;PHF6;RUNX1;SF3B1; CBL |
| 21.(1891) | CMML1 |  |  | DNMT3A;ROBO2;TET2 |
|  | AML | 14 | **CBL/11/c.1564-13C>T(56)**<br>**CEBPA/1/A265fs(37)K90fs(52)** | DNMT3A;ROBO2;TET2*2;CBL;CEBPA*2;EZH2 |
| 22.(4410) | RAEB1 |  | PTPN11/3/D61H(33) | EZH2; PTPN11;SETBP1;TET2 |
|  | AML | 12 | **PTPN11/3/D61H(48)** | EZH2; PTPN11;SETBP1;TET2 |
| 23.(2638) | CMML1 |  | PTPN11/14/P563R(43) | PTPN11; ROBO1; |
|  | AML | 16 | **PTPN11/14//P563R(45) /3/D61V(37)** | PTPN11x2; ROBO1;SF3B1 |
| 24(4133) | RAEB1 |  | CBL/9/R420Q(71)<br>TP53/4/c.306-2A>G(96) | ASXL1;CBL;ROBO2;TP53 |
|  | AML | 3 | CBL/9/R420Q(39)<br>TP53/4/c.306-2A>G(97)<br>**PTPN11/13/G507V(28)** | ASXL1;CBL;ROBO2;TP53;PTPN11 |
| 25.(3891) | RCMD |  | PTPRD/18/I930V ( 51 ) | ASXL1;PTPRD |
|  | AML | 8 | **PTPRD/18/I930V(46)** | ASXL1;PTPRD |
| 26.(702) | RCMD |  |  | ANKRD11;SRSF2 |
|  | AML | 28 | MPL/10/W515L(17)<br>**GATA2/6/R396W(15)** | ANKRD11;SRSF2 ; GATA2;IDH1;MPL |
| Myeloid Transcriptors |  |  |  |  |
| 27 (1931) | RAEB1 |  | **CEBPA/1/S9F(67)** | CEBPA;DNMT3A;TET2;U2AF1 |
|  | AML | 9 | **CEBPA/1/S9F(46)** | CEBPA;DNMT3A;TET2;U2AF1 |
| 28.(1243) | RCMD |  |  | ASXL1; U2AF1 |
|  | AML | 18 | **CEBPA/1/7_15del ( 40 )** | ASXL1; U2AF1; CEBPA; IDH1 |
| 29.(3288) | RN |  |  | KIF20B ; STAG2 ; TET2*2 |
|  | AML | 12 | **CEBPA/1/L338P(36)** | KIF20B ; STAG2 ; TET2*2 ; ASXL1 , CEBPA ; EZH2 |
| 30.(1090) | RARS |  | RUNX1/9/D344G(45 ) | DNMT3A ; DHX9 ; RUNX1 ; SRSF2 ; |
|  | AML | 11 | RUNX1/9/D344G(48)/7/R232W(20)<br>**CEBPA/1/Y285fs(32)** | DNMT3A ;DHX9 ;RUNX1*2 ;SRSF2 ; ASXL1 ;CEBPA ;SETBP1 |
| 31.(1018) | RAEB2 |  |  | DNMT3A; EZH2; TET2 |
|  | AML | 4 | FLT3:/9/D358V(53)<br>**CEBPA/1/A295V(39)** | DNMT3A; EZH2; TET2;BCOR;CEBPA ; FLT3 |
| 32.(3552) | RARS |  |  | SF3B1; TET2 |
|  | AML | 28 | **CEBPA/1/c238delineCSG(18)** | SF3B1; TET2; CEBPA;; IDH2;SRSF2; STAG2 |
| 33(3826) | RCMD |  |  | ASXL1;EZH2 |
|  | AML | 22 | **CEBPA/1/Y166X(48)P58fs(47)** | ASXL1;EZH2;CEBPA*2 |
| 34.(4196) | RCMD |  | PHF6/4/G93S(27)<br>RUNX1/1/R80C(51) | PHF6; RUNX1;SRSF2; |
|  | AML | 8 | PHF6/4/G93S(45)<br>RUNX1/1/R80C(78)<br>NRAS/2/G12D ( 34 )<br>**CEBPA/1/69_70del(11)** | PHF6; RUNX1;SRSF2; CEBPA;NRAS |
| 35.(4634) | RAEB2 |  | NPM1/7/L160fs(24)<br>NRAS/2/G13D(16) | NPM1 ; NRAS ; SF3B1;TET2 |
|  | AML | 2 | NPM1/7/L160fs(35)<br>NRAS/2/G13D(37)<br>**CEBPA/1/69_70del(9.8)** | NPM1 ; NRAS ; SF3B1;TET2 ; CEBPA |
| 36.(3536) | RCMD-RS |  | RUNX1/4/H105Y ( 40 ) | ASXL1;ROBO1;RUNX1;SF3B1;UPF3A |
|  | AML | 3 | **RUNX1/4/H105Y ( 34 )** | ASXL1;ROBO1;RUNX1;SF3B1;UPF3A |
| 37.(1350) | RAEB1 |  | PHF6/9/Y303X(56)<br>RUNX1/9/P333fs(29) | EZH2;PHF6;RUNX1;TET2 |
|  | AML | 2 | PHF6/9/Y303X(46)<br>**RUNX1/9/P333fs(17)** | EZH2;PHF6;RUNX1;TET2 |
| 38.(3479) | RCMD |  | RUNX1/3/R174Q(28) | ASXL1;RUNX1;TET2; ZRSR2 |
|  | AML | 15 | **RUNX1/3/R174Q(42)** | ASXL1;RUNX1;TET2*2;ZRSR2 ; |
| 39.(4143) | RAEB1 |  | RUNX1/4/S229fs ( 34 ) | BCOR;DNMT3A;IDH2;ROBO1;RUNX1 |
|  | AML | 3 | **RUNX1/4/S229fs ( 39 )** | BCOR;DNMT3A;IDH2;ROBO1;RUNX1 |
| 40.(1582) | CMML1 |  | PTPRD/22/c.962-9C>G(47) | KIF20B;PTPRD;SF3BA;TET2 |
|  | AML | 13 | PTPRD/22/c.962-9C>G(43)<br>***RUNX1/9/P340fs(40)*** | KIF20B;PTPRD;SF3BA;TET2;RUNX1 |
| 41.(4390) | RCMD |  | ETV6/7/M389R(33)<br>PHF6/10/R342X(27) | ASXL1;BCOR;ETV6;PHF6 |
|  | AML | 15 | PHF6/10/R342X(45)<br>***ETV6/7/M389R(42)*** | BCOR;ETV6;PHF6 |
| 42.(811) | RCMD |  |  | None |
|  | AML | 26 | ***ETV6/6/V345L ( 52 )***<br>***MPL/10/W515R(50)*** | ETV6;MPL |
| Tumor suppressors |  |  |  |  |
| 43.(1445) | RAEB2 |  | TP53/8/E287D(56) | ROBO1;TP53;UPF3A |
|  | AML | 27 | ***TP53/8/E287D ( 48 )*** | ROBO1;TP53;UPF3A |
| 44.(1694) | RAEB2 |  | TP53/7/M237I(74) | DNMT3A; TP53 |
|  | AML | 3 | ***TP53/7/M237I(80)*** | DNMT3A; TP53 |
| 45.(3880) | CMML1 |  | TP53/2/S83G(84) | TP53 |
|  | AML | 13 | ***TP53/2/S83G(60)*** | TP53 |
| 46(3678) | RAEB1 |  | TP53/5/530-535del (34) | TP53 |
|  | AML | 15 | ***TP53/5/530-535del ( 53 )*** | TP53 |
| 47.(1831) | RAEB1 |  | TP53/4/W91X(30) | ANKRD11;DNMT3A;TP53 |
|  | AML | 13 | ***TP53/4/W91X(30)/5/R158L(31)*** | ANKRD11;DNMT3A;TP53*2;KIF20B;STAG2 |
| 48.(2666) | RAEB1 |  | PTPRD/27/G809V(56) | DNMT3A;IDH2;PTPRD |
|  | AML | 18 | PTPRD/27/G809V(51)<br>**TP53/5/P151S(25)** | DNMT3A;IDH2;PTPRD;TP53; |
| 49.(1354) | RAEB1 |  |  | None |
|  | AML | 7 | **TP53/8/E287fs(40)/6/G199E(28)** | ASXL1 ; TP53x2 |
| 50.(2296) | RAEB2 |  |  | DHX9; ROBO1; |
|  | AML | 4.5 | **TP53/9/R303fs(85)** | DHX9;ROBO1; TP53 |
| 51.(843) | RCMD |  |  | SRSF2; TET2 |
|  | AML | 3 | **TP53/5/C135X(18)** | SRSF2;TET2;TP53 |
| 52.(1849) | RCMD |  | WT1/7/H428fs ( 31 ) | DNMT3A;IDH1;TET2; WT1 |
|  | AML | 8 | **WT1/7/H428fs ( 40 )** | DNMT3A;IDH1;TET2*2;WT1 |
| 53..(1670) | RCMD |  | GATA2/5/M388fs(41)<br>TP53/2/E11Q(51)<br>ETV6/5/V285M(49)<br>WT1/1/A115fs(12) | BCOR;ETV6;GATA2;TET2;TP53;U2AF1 ; WT1 |
|  | AML | 2 | GATA2/5/M388fs(40)<br>TP53/2/E11Q(50)<br>ETV6/5/V285M(53)<br>**WT1/1/A115fs(25)** | BCOR;ETV6;GATA2;TET2;TP53;U2AF1 ; WT1 |
| 54(3567) | RAEB2 |  | FLT3/12/c.1419-3->T(15)<br>NPM1/10/R262fs(43)/10W259delinsW<br>Q(44)/10/R262S(44)/7/W163L(42) | DHX9;DNMT3A;ANKRD11;FLT3; NPM*4; |
|  | AML | 4 | NPM1/10/R262fs(12)/10W259delinsW<br>Q(25)/10/R262S(19)/7/W163L(20)<br>**WT1/7/S169fs(32)** | DHX9;DNMT3A;ANKRD11; NPMx4;ROBO1;WT1 |
| 55.(1892) | RAEB2 |  | NPM1/11/L287f ( 35 ) | DNMT3A;IDH1;NPM1 |
|  | AML | 9 | **NPM1/11/L287f ( 31 )** | DNMT3A;IDH1;NPM1 |
| *The patients in whom last gene events could not be presumed by paired target sequencing analysis* |  |  |  |  |
| 56(1075) | RA |  |  | DHX9;IDH2;ROBO2; STAG2;UPF3A |
|  | AML | 20 |  | DHX9;IDH2;ROBO2; STAG2;UPF3A;TET2 |
| 57(1702)★ | RAEB2 |  |  | SRSF2 |
|  | AML | 19 |  | SRSF2 |
| 58.(1601)<br>★ | RAEB2 |  |  | SETBP1 |
|  | AML | 7.5 |  | SETBP1 |
| 59.(3831)<br>★ | RAEB2 |  |  | ANKRD11;IDH2 |
|  | AML | 4 |  | ANKRD11;IDH2 |
| 60.(3155)<br>★ | RCMD |  |  | ROBO1 |
|  | AML | 7.5 |  | ROBO1 |
| 61.(1619) | RAEB2 |  |  | DHX9;DNMT3A;ITIH3*2 |
|  | AML | 25 |  | DHX9;DNMT3A;ITIH3*2 |
| 62.(3437) | RA |  |  | DNMT3A;ROBO1 |
|  | AML | 3 |  | DNMT3A;ROBO1 |
| 63.(3053)<br>★ | RAEB2 |  |  | STAG2(11);UPF3 |
|  | AML | 40 |  | SETBP1(18);SRSF2 |
| 64.(4145) | RCMD |  |  | EZH2;IDH1 |
|  | AML | 14 |  | ASXL1;TET2*2 ; ZRSR2 |

Notes:1. The fourth column indicates the possible last event mutations. The mutations with italic and bold font were shown as the final presumed AML transformation-related mutations (TRM) according to the definition in the METHOD part. 2. The paired target sequencing analysis from the last nine patients could not define the TRM, so five (with star signals)of them accepted the whole exome sequencing to look for possible TRM which were not included in 39 target genes.

1. Transformation-related mutations (TRM) were identified in 55 of 64 patients (85.9%), according to the steps described in METHODS.
2. Among these 55, TRM were detected only at AML transformation in 35 patients (representing 63.6% of these cases). In the 20 remaining cases, these mutations were detected at the point of diagnosis (Table 1). Patients with TRM in only their sAML sample trended toward a longer time to transformation (mean 13.4 months vs 9.6 months for the first; *P*=0.106) and no difference in lower risk (<RAEB2/CMML2) cases at diagnosis (80.0% vs 75.0% for the second; *P*=0.431).
3. From Table 1, 2 and Figure 1, the following defined TRMs were present in descending order of frequency. First were active signaling mutations:*KRAS*/*NRAS* in 11 patients (two also had *CBL* or *PTPN11* as TRM); *FLT3* in six patients (one also had CBL); *CBL* in six patients (three had *NRAS*; *FLT3* and *CEBPA* respectively); and *PTPN11* in four (one had *NRAS*); the remaining two had *PTPRD* and *GATA2*, respectively. The next set of mutations comprised myeloid transcription factors: *CEBPA* in 10 patients (one had *CBL*); *RUNX1* in five patients, and *ETV6* in two patients. The next set of mutations comprised tumor suppressor genes: *TP53* mutations in nine patients; *WT1* mutations in three patients, and *NPM1* in one patient. *KIT* mutations were not detected in this group of patients. The TRM events seemed to be highly enriched in about 10 genes. Figure 1 shows the pattern of these mutation events, demonstrating that the involvement style of *FLT3* was not the same as that of FLT3-ITD, which is more often observed in *de novo* AML.
4. In 6 of these 55 paired samples with candidate transformation events, we sequenced an additional sample between MDS diagnosis and AML transformation (Figure 2). Of these, 3 samples (UPN1243, 3288, and 4390) showed evidence of the TRM emerging before phenotype change, i.e., the TRM event emerged while the disease was still in the MDS stage, with sAML transformation occurring soon thereafter.
5. Analysis of changing mutation profiles and VAF, as well as the logical relations between them, revealed differing patterns of evolution from MDS to sAML. Of the 55 patients with candidate mutation events, most (25 cases) demonstrated AML-transformation-related progression by linear evolution from the founding clone (Figure 3a). Five cases showed linear evolution from subclones (Figure 3b). Only four cases demonstrated sweeping clonal evolution (Figure 3c). The evolution pattern of the remaining 21 patients either could not be defined by changes in mutations or VAF (4 cases), or harbored no candidate alterations before or after progression to AML.
6. Rivalry clonal evolution patterns were observed. Some original TRM were replaced or outgrown by other mutations at sAML transformation, such as in patients 2 (UPN3430), 5 (UPN809); 9 (UPN4603), 13 (UPN2570), and 54 (UPN3567) (Table 1)
7. Although differences existed among TRM, TET2/RUNX1/ASXL1 /DNMT3A /EZH2/U2AF1 mutations were the most common initial/driver mutations for these AML-transformed patients (Table2). However, TET2/ASXL1/BCOR /ROBO1 mutations were most commonly accompanied by those transformation-related events when transformation occurred (Table2).

**Table 2.** Characteristics summary for the detailed data from Table 1.

| Last event mutation | cases | Emerged at sAML(n) | Primary diagnosis≥ RAEB2/CMML2(n) | Most common earlier mutations at MDS diagnosis(top 2)(top4 in Total)(n) | Most common partner mutations at transformation(top 1)(top2 in Total)(n) |
| --- | --- | --- | --- | --- | --- |
| ***NRAS/KRAS*** | 11(2) | 9/11 | 3/11 | RUNX1(8)/ASXL1(5)TET2(5)/EZH2(4)<br>STAG2(4) | BCOR(3) |
| ***CEBPA*** | 10(1) | 9/10 | 2/10 | TET2(6)/DNMT3A(4)/ASXL1(2)EZH2(2)<br>SF3B1(2)RUNX1(2)SRSF2(2)U2AF1(2)) | ASXL1(2)EZH2(2) |
| ***TP53*** | 9 | 5/9 | 3/9 | DNMT3A(3)/ROBO1(2) | ASXL1(1)KIF20B(1)Stag2(1) |
| ***FLT3*** | 6(1) | 5/6 | 0/6 | U2AF1(3)/IDH2(2)RUNX1(2)SRSF2(2) | ROBO1(2)TET2(2) |
| ***CBL*** | 6(3) | 4/6 | 2/6 | U2AF1(3)/ANKRD11(2)ASXL1(2)DNMT3<br>A(2)<br>STAG2(2) | TET2(2) |
| ***RUNX1*** | 5 | 1/5 | 0/5 | TET2(3)/ASXL1(2)SF3B1(2) | none |
| ***PTPN11*** | 4(1) | 3/4 | 0/4 | EZH2(2)SETBP1(2) | SF3B1(1) |
| ***WT1*** | 3 | 1/3 | 1/3 | DNMT3A(2)TET2(2) | ROBO1(1) |
| ***Total*** | 54(4) | 37/54 | 11/54 | TET2(16)/RUNX1(12)/ASXL1(11)DNMT3<br>A(11)/EZH2(8)/U2AF1(8) | TET2(4)/ASXL1(3)BCOR(3)ROBO1(3) |

NOTE: 1.The data from the cases with less observed TRM (less than 2 cases in total) is not included in this Table. 2. The number in the brackets of the second column represented the cases who had another presumed TRM besides the mainly presumed TRM. 3. The mutations listed in the last column means the mutations which emerged together with the presumed TRM, may playing roles during the MDS-sAML transformation.

**Figure 1.**
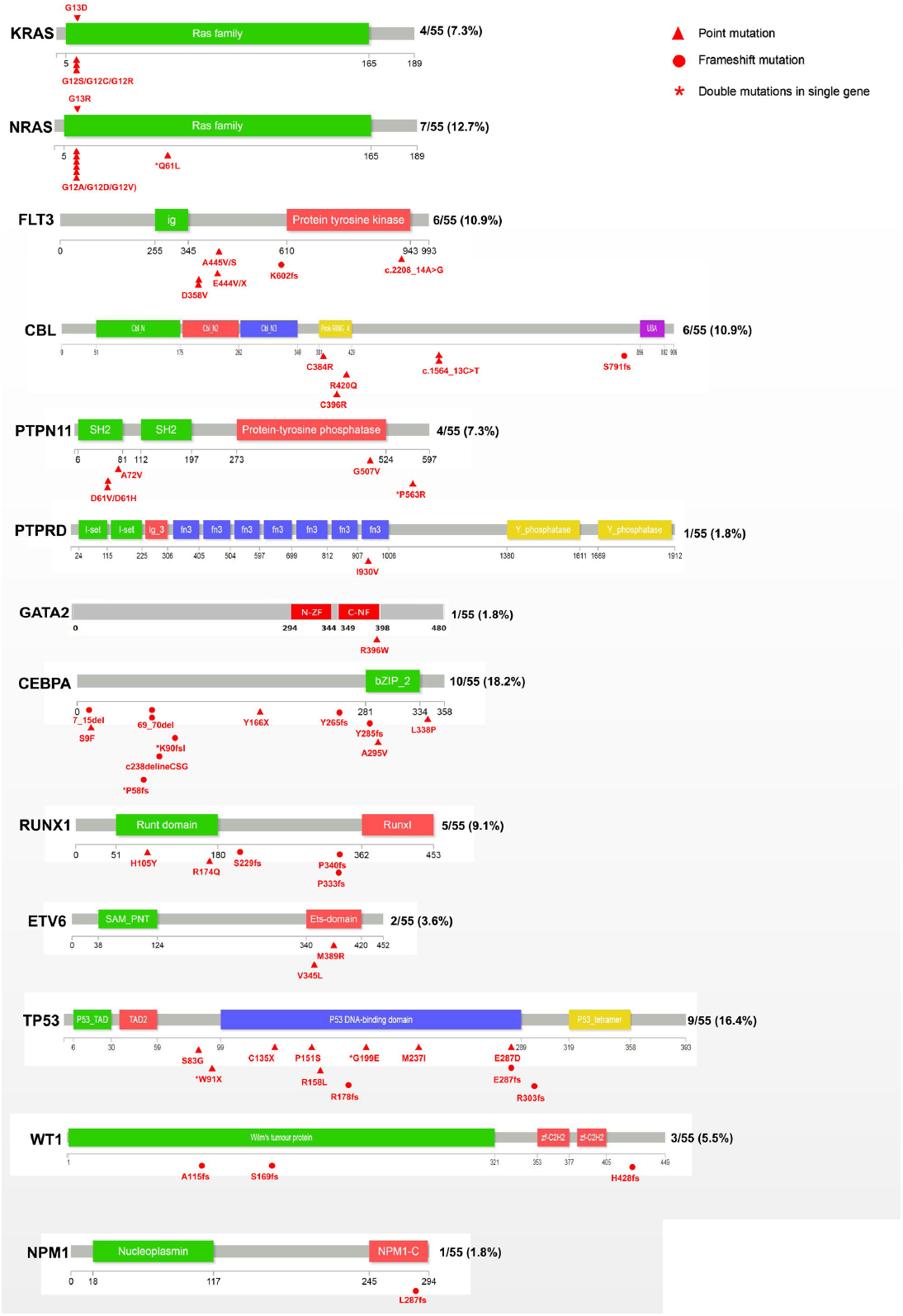
Occurrence frequency and sites of last mutation events. All mutations, including types and sites, are shown for 13 AML transformation-related mutations, according to the definitions. Triangles and circles indicate point and frameshift mutations, respectively. Stars indicates double mutations occurring in a single gene.

**Figure 2.**
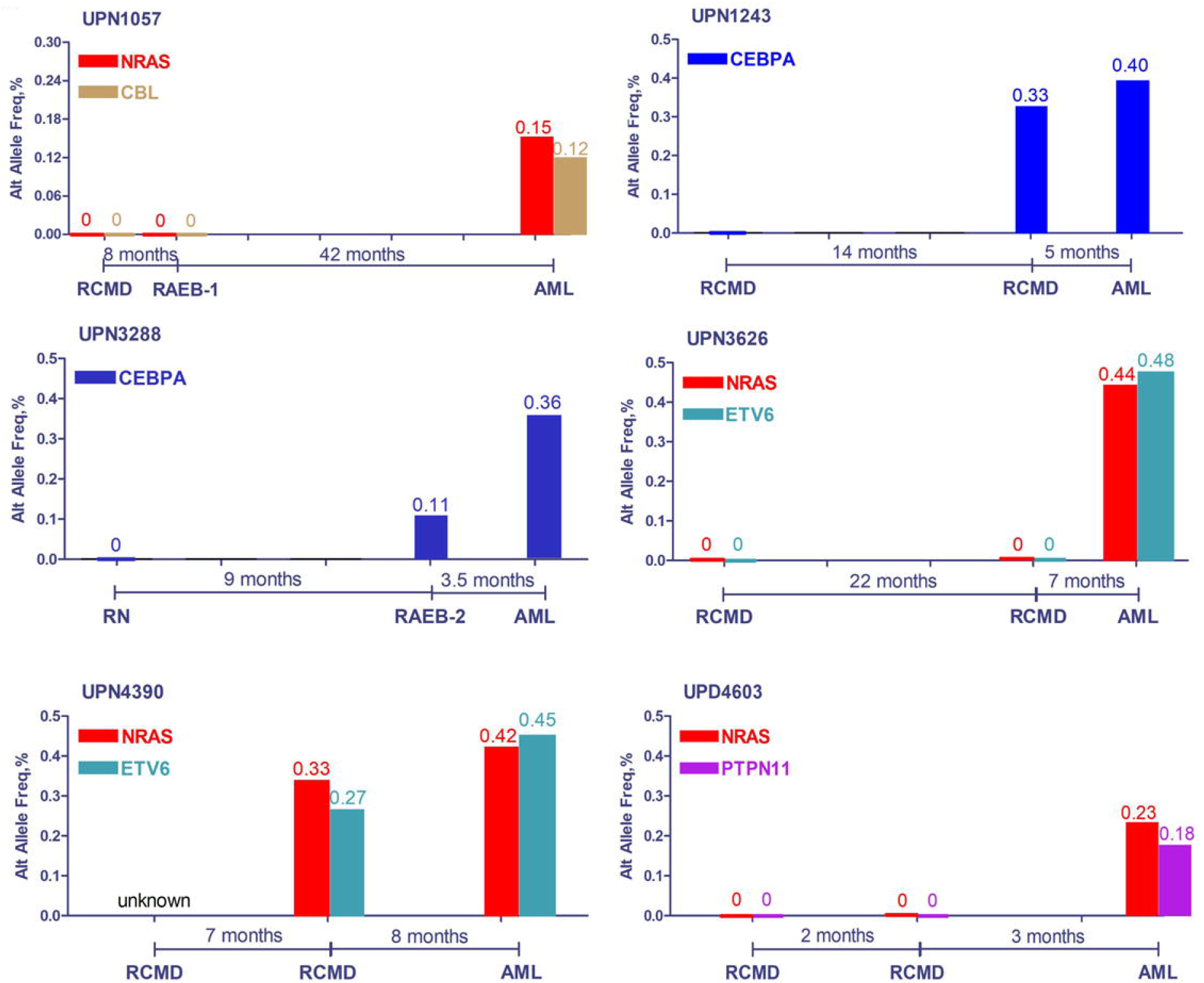
Evolution route for six cases that were analyzed by an additional sequencing assay between MDS diagnosis and AML transformation.

**Figure 3.**
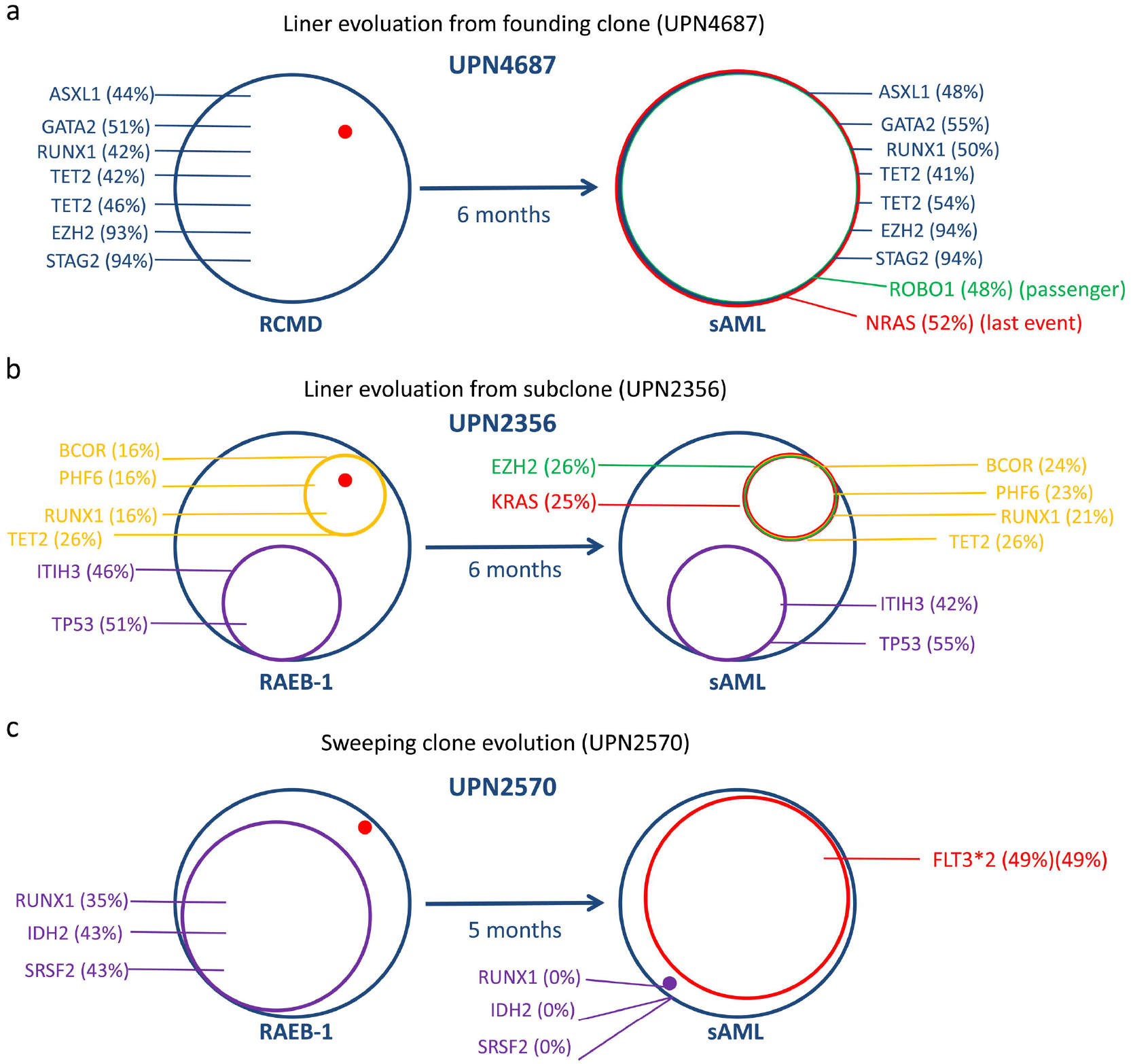
Three evolution patterns for AML transformation. **a**. Linear evolution, based on the founding clone, is shown for UPN4687. Similar cases include UPN809, 1057, 3435, 3626, 1792, 3183, 1891, 1243, 3288, 1090, 1018, 3552, 3826, 4196, 4634, 1582, 1831, 2296, 843, 702, 3430, 3541, 2638, and 4133. **b**. Linear evolution from a subclone is shown for UPN2356. Similar cases include UPN4603, 2577, 692, and 2666. **c**. Sweeping clone evolution is shown for UPN2570. Similar cases include UPN811, 1354, and 3567. [Number] indicates VAF; [H] indicates homozygous mutations; red indicates presumed last mutation; green indicates newly emerged partner mutations, which could not be presumed to be AML transformation-related mutations.

### Whole exome sequencing

Samples from five of the nine patients whose target sequencing showed no TRM but for whom sufficient DNA extract was still available were further subjected to whole exome sequencing (WES) (patients starred in Table 1). Figure 4 presents the results of WES from these cases. In addition to MDS-related gene mutations, each gained at least one TRM (involving activated signaling, transcription factor, or tumor suppressor genes) at the AML transformation point. Specifically, UPN1601 and UPN 3053 acquired NOTCH1 and PAK5 mutations, respectively (both involved in signaling transduction), during disease progression. NOTCH1 mutations are reported to occur frequently in acute leukemia [14], whereas PAK5 contributes to the proliferation of tumor cells as serine/threonine protein kinase [15]. UPN1702 and UPN3831 acquired MLLT10 and NCOR2 mutations, respectively (both involved in transcription regulation), at the AML stage. MLLT10 is a histone lysine methyltransferase, which participates in AML pathogenesis via the formation of fusion gene MLLT10-MLL [16]. NCOR2 regulates gene transcription as a part of a histone deacetylases complex, and its dysfunction leads to functional abnormality in hematopoietic stem cells [17]. UPN3155 acquired a STK11 mutation (a tumor suppressor) after disease progression. It has been reported that STK11 regulates cell polarity and functions as a tumor suppressor [18].

**Figure 4.**
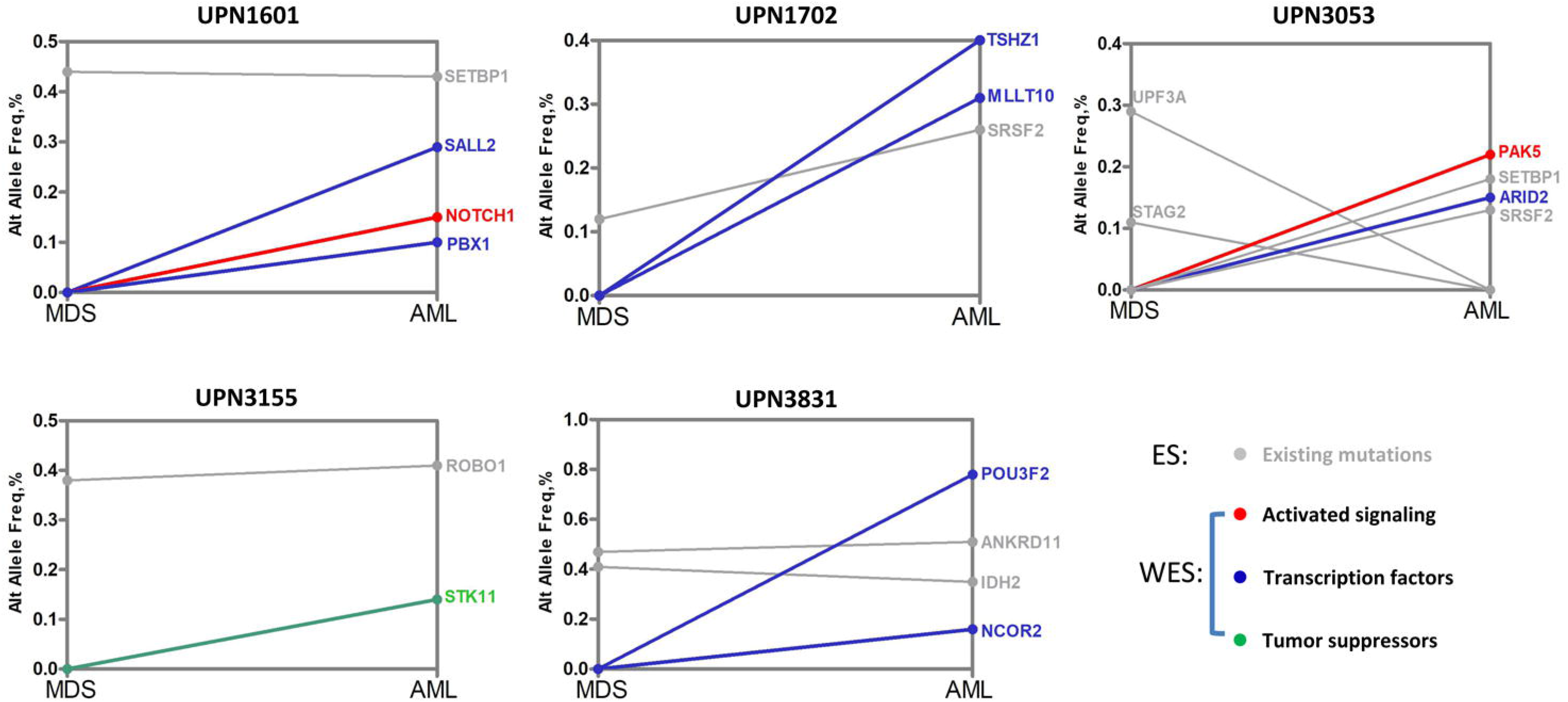
Whole exome sequencing revealed other atypical last mutation events. Five patients showed negative results for typical AML transformation-related mutations. However, whole exome sequencing revealed some other atypical AML transformation-related mutations. Gray indicates existing mutations; red indicates activated signaling; blue indicates transcription factors; green indicates tumor suppressors.

### Single-cell RNA sequencing

As mentioned above, abnormal cell signaling induced by gene mutation, such as RAS genes or *PTPN11* may be critical for the transformation of MDS into AML. However, it is still unclear whether a RAS mutation is a requirement to activate RAS signaling pathways. To explore this question, we used single-cell RNA transcription sequencing to study the association between RAS mutation and RAS signaling in UPN4674 and UPN4763 (before and after disease progression). Two s (*NRAS* and *PTPN11* mutations) occurring during the AML stage are core genes in RAS signaling. As shown in Figure 5a and 5b, this patient presented several abnormal cell types (orange, UPN4674; blue, UPN4763 in Figure 5a). Gene classification analysis showed that aberrant granular-mononuclear progenitors (GMP), common myeloid progenitor (CMP), megakaryocyte-erythroid progenitor (MEP) and monocytes (Mono) were present during the AML stage (Figure 5b). We focused on differences in the GMP population were RAS signaling. Integrated analysis based on single-cell sequencing indicated that several RAS signaling related-genes are expressed at high levels in GMP after disease progression (Figure 5c). These genes have been reported to participate in the activation of RAS signaling [19]. Similarly, RAS signaling related-genes such as *FLT3, INSR*, and *CDC42* are also expressed at high levels in MEP, CMP, and Mono groups after disease progression (Figure 5d). These genes are closely associated with cell proliferation. Interestingly, apoptosis-related gene *BAD* is down-regulated in GMP, MEP, CMP, and Mono groups after disease progression. These data suggest that transformation-related mutations gene mutations induce the redistribution of clonal cells (leading to an increased number of morphological blasts and monocytes) via the activation of cell signaling, and further leads to AML transformation.

**Figure 5.**
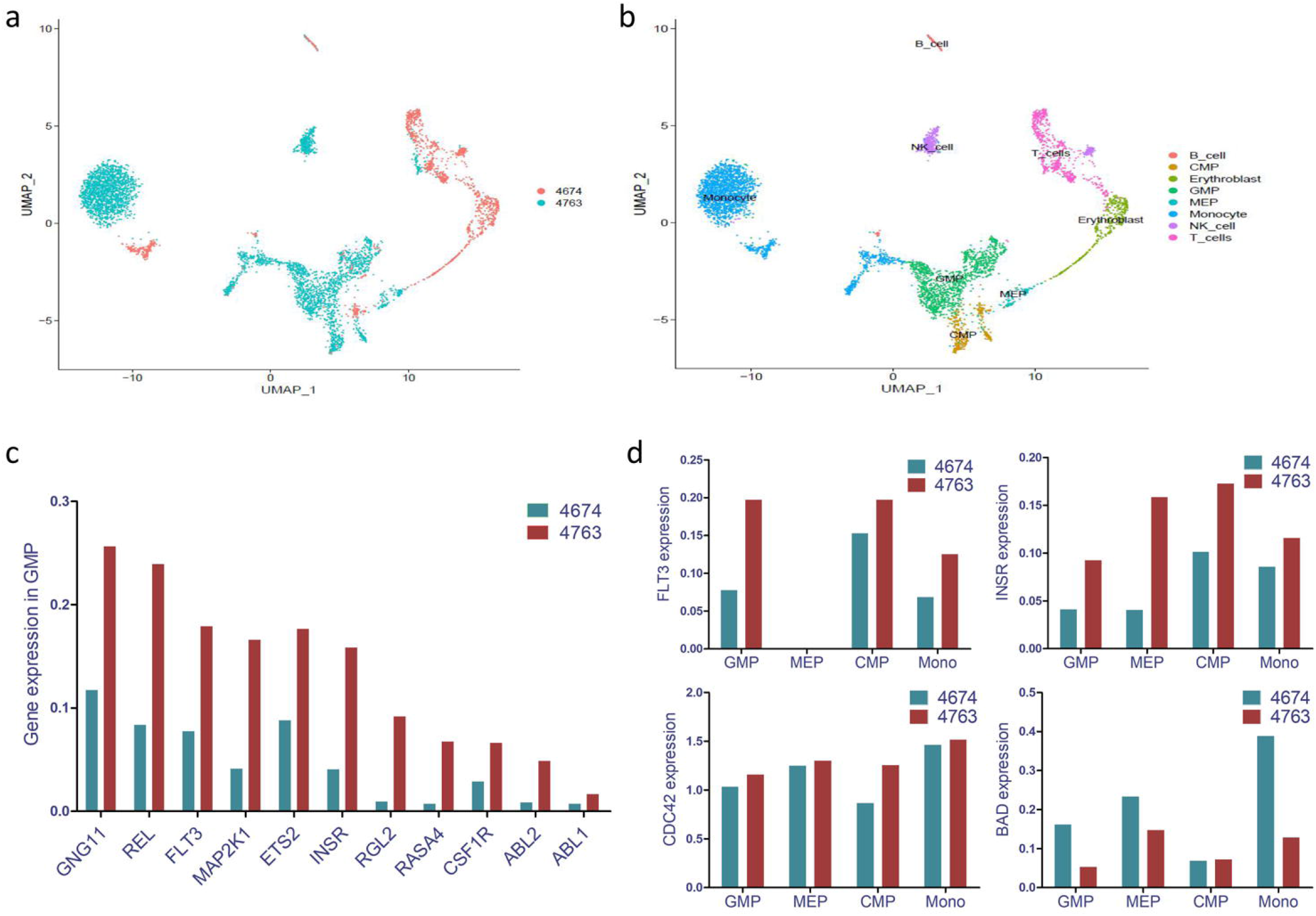
Single-cell RNA transcription group sequencing revealed the linear association of gene mutations with activation of targeted signaling. (**a**) Patient presented with several abnormal cell groups (blue) after disease progression. (**b**) Cluster analysis suggested that abnormal GMP, CMP, MEP, and monocytes increased in the AML stage. (**c**) Integrated analysis indicated that several RAS signaling related-genes were highly expressed in GMP after disease progression. (**d**) RAS signaling related-genes, such as FLT3, INSR, and CDC42 were also highly expressed in MEP, CMP, and Mono groups. However, apoptosis-related gene BAD is down-regulated in GMP, MEP, and CMP.

## Discussion

The pathogenesis of sAML differs from that of *de novo* AML in several respects including clinical progression and prognosis [20]. Despite poor response to contemporary therapies, including stem-cell transplantion, MDS-derived AML exhibits a relatively long pre-transformation duration, and occurs only in about one third of all MDS [7]. If this transformation requires triggering factors (defined as TRM), and these factors are robust biomarkers of progression, it may be possible to develop counter mechanisms to prevent transformation.

In this study, we sequenced 64 paired bone marrow DNA samples, first at MDS diagnosis and then immediately after AML transformation. Fifty-five out of the 64 cases (85.9%) were positive for TRM, i.e., events involving active signaling, transcription factors, and tumor suppressors. Sequencing of 39 target genes could not detect any TRM in 9 out of the 64 cases. Of these, paired whole exome sequencing was performed for 5 cases for which sufficient DNA could be extracted from BM samples. Each of these patients carried at least one TRM outside of the 39 target genes, involving active signaling, myeloid transcription, or tumor suppressor genes (Figure 4). No patient with MDS who developed sAML did so only by acquiring mutations involved in epigenetic modulation, RNA splicing, or cohesins.. This finding suggests that the existence or emergence of certain last event-like mutations may be considered a prerequisite for MDS to transform into sAML, rather than just occasional or accompanying events [9,10].

Among the 60 patients with detected TRM, in 40 patients (66.7%; including 35 by targeted sequencing and 5 by WES) the TRM appeared to emerge at the point of AML transformation, and the remaining 20 patients harbored putative last mutations at the time of MDS diagnosis. Then it is necessary to tell which the original is? The existence/emerge of TRM? Or the AML transformation itself? We considered that the pre-existence or emergence of TRM should be the reason for AML transformation. First, 20/60 (33.3%) cases harbored TRM at MDS diagnosis; The duration of transformation was moderately shorter in these cases than in those where the TRM were only identified at the transformation point (mean 9.8 vs. 13.4 months). Second, due to the relatively longer corresponding intermittence between time points at diagnosis and AML transformation for paired sequencing, some important information may have been missed. When additional time points were added, supportive data were obtained from some patients (UPN1243, 3288, and 4390 in Figure 2). When TRM emerged in these patients, their disease was still at the low-risk or pre-AML stage, but transformed to AML quickly. Finally, for one patient (UPN 4603), immediately after the occurrence of the AML phenotype, accompanied by NRAS and PTPN11 mutations, single-cell RNA sequencing showed both active RNA transcription and activation of the RAS/PTPN11 signaling pathway [21,22]. Given that the gene mutations occurred prior to RNA/protein transcription/translation, pathway activation, and ultimately, phenotype alteration. These results support our findings that TRM and activated RAS signaling can precede phenotypic transformation to AML.

Most of the TRM clones showed linear evolution from the founding clones (25 cases, compared to 5 cases with linear evolution in a pre-existing subclone; and 4 cases by clone sweeping) (Figure 3). This is somewhat different from published data [23,24], may because of that a part of patients did not presented clonal progressing process, such as those 20 cases whose TRM pre-existed at MDS diagnosis. In addition, we detected common pre-existing mutations involving *TET2*/*RUNX1*/*ASXL1*/*DNMT3A* (Table 2), indicating that these mutations represent precursor mutations for these AML cases. As for *TET2*/*ASXL1*/*BCOR*/ *ROBO1* mutations, they emerged as common partners of TRM (Table 2), possibly playing some roles in the transformation process. Some mutated genes such as RUNX1 (highly prevalent among patients with chromosome 7 abnormalities) and BCOR (common in chromosome normal cases) [25] could be responsible either for the MDS phenotype or driving AML progression. These dynamic genetic features contribute to the complexity and heterogeneity of this disease and may become a key to deciphering the rule for sAML occurrence.

Of note, the majority of detected TRM were from a cluster of 10 related genes. *KIT* mutation was never detected in this group of patients, and NPM1 was only detected in one case. The latter could be attributed to the favorable responses of the MDS patients who harbor NPM1 mutations towards HMAs (decitabine), thus blocking sAML transformation [26]. Together with *TP53*, mutations in *NRAS*/*KRAS, CEBPA*, and *FLT3* were observed in 63.6% of the 55 cases for which TRM were detected by targeted sequencing. When *RUNX1* /*CBL*/*PTPN11*/*WT1* and *ETV6* were included, only 10 mutated genes accounted for almost all the patients with MDS (94.5%) who experienced AML progression. In view of the rapid development of mutation-specific targeted therapy, we can hope to pharmacological block AML transformation in MDS patients at high transformation risk with regular monitoring for last mutations and administering effective corresponding targeted therapy.

In summary, somatic mutations in activated signaling, transcription factors, or tumor suppressor genes appear to be a precondition for AML transformation in myelodysplastic syndromes. The high propensity to acquire transformation-related gene mutations is worthy of further research as targets to be exploited with novel therapies.

## Acknowledgments

This work was supported by the National Natural Science Foundation of China (81770120 and 81770122). We thank Shanghai Tianhao Inc. for providing assistance in exome sequencing and data analysis. We sincerely thank Dr. Rafael Bejar for providing constructive suggestion and revision.

## Conflict of Interest

There are no financial and non-financial interests to declare.

## Author contributions

X.L. was the principal investigators who conceived the study. C.-K.C., F.X., and L.-Y.W. carried out most of the experiments. L.-X.S.. and Q.H. were responsible for bioinformatics investigation. J.G. and Y.T. participated in the preparation of biological samples. Z.Z., D.W., L.-Y.Z., C.X. and J.-Y.S. helped gather detailed clinical information for the study and helped to carry out clinical analysis. X.L.,C.-K.C and F.X. wrote the manuscript.

## Notes

This study was supported by the National Natural Science Foundation of China (grant nos. 81770120 and 81770122).

### Competing Interest Statement

The authors have declared no competing interest.

### Summary of Updates

author numbers updated,and manuscript updated

## References

1. Vetrie D, Helgason GV, Copland M. The leukaemia stem cell: similarities, differences and clinical prospects in CML and AML. Nat Rev Cancer. 2020 Mar;20(3):158–173.

2. de Thé H, Chen Z. Acute promyelocytic leukaemia: novel insights into the mechanisms of cure. Nat Rev Cancer. 2010 Nov;10(11):775–83.

3. Lin S, Mulloy JC, Goyama S. RUNX1-ETO Leukemia. Adv Exp Med Biol. 2017;962:151–173.

4. Douet-Guilbert N, Chauveau A, Gueganic N, Guillerm G, Tous C, Le Bris MJ, Basinko A, Morel F, Ugo V, De Braekeleer M. Acute myeloid leukaemia (FAB AML-M4Eo) with cryptic insertion of cbfb resulting in cbfb-Myh11 fusion. Hematol Oncol. 2017 Sep;35(3):385–389.

5. Hasserjian RP. Acute myeloid leukemia: advances in diagnosis and classification. Int J Lab Hematol. 2013 Jun;35(3):358–66.

6. Park S, Cho BS, Kim HJ. New agents in acute myeloid leukemia (AML). Blood Res. 2020 Jul 31;55(S1):S14–S18.

7. Sperling AS, Gibson CJ, Ebert BL. The genetics of myelodysplastic syndrome: from clonal haematopoiesis to secondary leukaemia. Nat Rev Cancer. 2017 Jan;17(1):5–19.

8. Raza A, Galili N. The genetic basis of phenotypic heterogeneity in myelodysplastic syndromes. Nat Rev Cancer. 2012 Dec;12(12):849–59.

9. Walter MJ, Shen D, Ding L, Shao J, Koboldt DC, Chen K, Larson DE, McLellan MD, Dooling D, Abbott R, Fulton R, Magrini V, Schmidt H, Kalicki-Veizer J, O’Laughlin M, Fan X, Grillot M, Witowski S, Heath S, Frater JL, Eades W, Tomasson M, Westervelt P, DiPersio JF, Link DC, Mardis ER, Ley TJ, Wilson RK, Graubert TA. Clonal architecture of secondary acute myeloid leukemia. N Engl J Med. 2012 Mar 22;366(12):1090–8.

10. Kim T, Tyndel MS, Kim HJ, Ahn JS, Choi SH, Park HJ, Kim YK, Yang DH, Lee JJ, Jung SH, Kim SY, Min YH, Cheong JW, Sohn SK, Moon JH, Choi M, Lee M, Zhang Z, Kim DDH. The clonal origins of leukemic progression of myelodysplasia. Leukemia. 2017 Sep;31(9):1928–1935.

11. Li X, Xu F, Wu LY, et al. A genetic development route analysis on MDS subset carrying initial epigenetic gene mutations. Sci Rep. 2020 Jan 21;10(1):826.

12. Vardiman JW, Thiele J, Arber DA, Brunning RD, Borowitz MJ, Porwit A, Harris NL, Le Beau MM, Hellström-Lindberg E, Tefferi A, Bloomfield CD. The 2008 revision of the World Health Organization (WHO) classification of myeloid neoplasms and acute leukemia: rationale and important changes. Blood. 2009; 114(5): 937–951.

13. Bennett JM, Catovsky D, Daniel MT, Flandrin G, Galton DA, Gralnick HR, Sultan C. Proposals for the classification of the myelodysplastic syndromes. Br J Haematol. 1982; 51:189–99.

14. Tohda S. NOTCH signaling roles in acute myeloid leukemia cell growth and interaction with other stemness-related signals. Anticancer Res. 2014 Nov;34(11):6259–64.

15. Li YK, Zou J, Ye DM, Zeng Y, Chen CY, Luo GF, Zeng X. Human p21-activated kinase 5 (PAK5) expression and potential mechanisms in relevant cancers: Basic and clinical perspectives for molecular cancer therapeutics. Life Sci. 2020 Jan 15;241:117113.

16. Van Limbergen H, Poppe B, Janssens A, De Bock R, De Paepe A, Noens L, Speleman F. Molecular cytogenetic analysis of 10;11 rearrangements in acute myeloid leukemia. Leukemia. 2002 Mar;16(3):344–51.

17. Yan M, Burel SA, Peterson LF, Kanbe E, Iwasaki H, Boyapati A, Hines R, Akashi K, Zhang DE. Deletion of an AML1-ETO C-terminal NcoR/SMRT-interacting region strongly induces leukemia development. Proc Natl Acad Sci U S A. 2004 Dec 7;101(49):17186–91.

18. Tarumoto Y, Lu B, Somerville TDD, Huang YH, Milazzo JP, Wu XS, Klingbeil O, El Demerdash O, Shi J, Vakoc CR. LKB1, Salt-Inducible Kinases, and MEF2C Are Linked Dependencies in Acute Myeloid Leukemia. Mol Cell. 2018 Mar 15;69(6):1017-1027.e6.

19. Li S, Balmain A, Counter CM. A model for RAS mutation patterns in cancers: finding the sweet spot. Nat Rev Cancer. 2018 Dec;18(12):767–777.

20. Winer ES. Secondary Acute Myeloid Leukemia: A Primary Challenge of Diagnosis and Treatment. Hematol Oncol Clin North Am. 2020 Apr;34(2):449–463.

21. Quan X, Deng J. Core binding factor acute myeloid leukemia: Advances in the heterogeneity of KIT, FLT3, and RAS mutations (Review).Mol Clin Oncol. 2020 Aug;13(2):95–100.

22. Liu X, Ye Q, Zhao XP, Zhang PB, Li S, Li RQ, Zhao XL. RAS mutations in acute myeloid leukaemia patients: A review and meta-analysis. Clin Chim Acta. 2019 Feb;489:254–260.

23. da Silva-Coelho P, Kroeze LI, Yoshida K, Koorenhof-Scheele TN, Knops R, van de Locht LT, de Graaf AO, Massop M, Sandmann S, Dugas M, Stevens-Kroef MJ, Cermak J, Shiraishi Y, Chiba K, Tanaka H, Miyano S, de Witte T, Blijlevens NMA, Muus P, Huls G, van der Reijden BA, Ogawa S, Jansen JH. Clonal evolution in myelodysplastic syndromes. Nat Commun. 2017 Apr 21;8:15099.

24. Makishima H, Yoshizato T, Yoshida K, Sekeres MA, Radivoyevitch T, Suzuki H, Przychodzen B, Nagata Y, Meggendorfer M, Sanada M, Okuno Y, Hirsch C, Kuzmanovic T, Sato Y, Sato-Otsubo A, LaFramboise T, Hosono N, Shiraishi Y, Chiba K, Haferlach C, Kern W, Tanaka H, Shiozawa Y, Gómez-Seguí I, Husseinzadeh HD, Thota S, Guinta KM, Dienes B, Nakamaki T, Miyawaki S, Saunthararajah Y, Chiba S, Miyano S, Shih LY, Haferlach T, Ogawa S, Maciejewski JP. Dynamics of clonal evolution in myelodysplastic syndromes.Nat Genet. 2017 Feb;49(2):204–212.

25. Xu F, Wu LY, He Q, et al. Exploration of the role of gene mutations in myelodysplastic syndromes through a sequencing design involving a small number of target genes. Sci Rep. 2017 Feb 21;7:43113.

26. Wu L, Li X, Xu F, et al. NPM1 mutation with DNMT3A wild type defines a subgroup of MDS with particularly favourable outcomes after decitabine therapy. Br J Haematol. 2020;189(5):982–984.

